# Genetic Algorithm with Rank Selection optimises robust parameter estimation for systems biology models

**DOI:** 10.1101/2022.02.22.481394

**Authors:** Gemma Douilhet, Mahesan Niranjan, Andres Vallejo, Kalum Clayton, James Davies, Sofia Sirvent, Jenny Pople, Michael R Ardern-Jones, Marta E Polak

## Abstract

The ability to reliably predict and infer cellular responses to environmental exposures would offer a major advance in the investigation of immune regulation in health and disease. One possible approach is the use of *in silico* modelling. Design of such a mathematical kinetic model would be based on existing knowledge of a biological system and utilise a partial data set to parameterise. However, the process of parameter estimation, key for the accuracy of the model, is difficult to conduct by hand, and thus a computational alternative is necessary. We report the utility of Genetic Algorithm with Rank Selection (GARS) as a parameter estimation tool on multiple biological models, including heat shock, signal transduction via ERK, circadian rhythm and NFκB systems, where it showed strong accuracy and superiority to the Extended Kalman Filter method, Algebraic Difference Equations, and MATLAB fminsearch approaches. GARS parameter estimation is a valuable tool for biological data because it reliably infers system behaviour from partial data sets, allowing for the prediction of cellular responses to environmental exposures.

## Introduction

Accurate regulation of cellular behaviour in response to signals from external environment and local tissue microenvironment is critical for health and homeostasis. However, this process is often aberrant in disease states, and during aging^1^. The ability to comprehensively analyse signalling events is key for identification of the drivers of pathology and offers the opportunity to predict signalling outcome and identify targets for interventions. Computational modelling offers the most promising way to approach the problem, providing the mathematical framework for modelling the resting state of signalling systems, including disease-specific steady states and predictive in silico testing of the effectiveness of therapeutic intervention ^2^, predicting the cell and system behaviour during prolonged exposure to signalling stimulus, and the outcome of multiple signalling events ^1,3^. By measuring cellular changes after manipulation of the stimulus *in vitro*, it is relatively easy to develop equations to predict outcomes from events which induce a simple linear signalling cascade. However, linear models cannot encompass complex systems with multiple pathways and feedback loops. Furthermore, in complex systems, multiple linear models cannot be developed to matrix because detailed measurement of each step of live intracellular signalling pathways in *in vitro* to inform the equations is currently not possible. Therefore, to achieve a predictive tool for complex systems, computational approaches using machine learning offer the most promising solution.

Constructing a predictive model of a biological system critically depends on correctness of several steps: reconstruction of network architecture, design of kinetic equations describing system dynamics, and derivation of kinetic parameters.^4–6^ Reconstruction of network architecture is usually based on prior knowledge of the biological system, capturing the known logic of interactions between biological entities in depicted in a diagram^4–7^. Stochastic modelling of distributed systems, using Michaelis-Menten kinetics based rate laws and mass action kinetic models have been successfully used for design of kinetic equations simulating small biological networks, and provided insights into the mechanisms regulating gene, signalling and metabolic regulatory network behaviour^8–10^. In contrast, estimation of kinetic parameters, such as rate constants and half-lives of molecular species, critical for correct representation of system dynamics, is a key challenge for mechanistic modelling of signal flow in a mathematical model of a network^11^. For *in silico* models, such parameters are usually derived from *in vitro* experiments or hand-tuned during model development, in a laborious and often inaccurate process. Advanced machine learning algorithms offers an alternative for reliable estimation of parameters and states, converging from initial distributions during system perturbations and infer the regulatory circuits from single-cell gene expression data^12,13^.

Currently, a number of computational methods exist for estimating parameter values from synthetic and experimental data sets. Lillacci and Khammash^14^, and Liu *et al*^13^ used the Extended Kalman Filter (EKF) as a form of parametric parameter estimation to determine parameter values for the heat shock model. Liu *et al* utilised a comprehensive synthetic data set of 1000 sample points, whereas Lillacci and Khammash sampled a dataset of only 13 data points^14^. Both papers added Gaussian noise to their synthetic data points. The EKF was also employed by Sun *et al* for the ERK Network model to estimate the 11 parameters using an training data from an experimental time course of 10 time points measuring each of the 11 proteins in the model^15^. However, the EKF requires an initial user-defined estimation for each parameter value, which is almost impossible when utilising a model for which there is little information available about the kinetic rates, and the value of which can have a profound effect on the resulting final parameter estimation. As a consequence of the parametric model making Gaussian assumption about its distribution, the EKF also has the additional complication of having no bounded search range, meaning that parameter value estimations can be below zero, as was the case for some values in the heat shock model^13^. These negative values may provide a close approximation to the true parameter values, however they are not biologically feasible, and they can therefore have a significant effect on the model fit to the data (Table S1), which means that accurate prediction using the model is unlikely. A similar problem is encountered when using Maximum Likelihood Estimation approaches such as MATLAB’s fminsearch program^13^. In addition, Liu et al found that when more than one parameter value were being estimated, using sequential methods such as the EKF was more effective than the Maximum Likelihood Estimations^13^.

As a potential solution to these issues, the genetic algorithm has been proposed as a tool for estimating parameter values to fit data sets from biological networks. However, data fitting alone is not sufficient and we must look towards utilising biological models and parameter estimations to predict unseen data.

Here we present a Genetic Algorithm with Rank Selection (GARS, Figure 1) as a tool for estimating parameter values which will not only provide a good model fit to both synthetic and experimental data, but can also be utilised to predict data after observations have ceased, or following changes to the biological system. This GARS requires no user-defined initial input, meaning that it can be employed on models with no prior knowledge of the system kinetic rates, and although it does require a given search space, we show that this can be widened with no negative impact on the model fit. We validated the GARS on a number of biological models to demonstrate its ability to be used with systems of different sizes and complexity, and established it as the most effective method of parameter estimation when compared to others that were previously utilised on the ERK network^15,16^.

**Figure 1:**
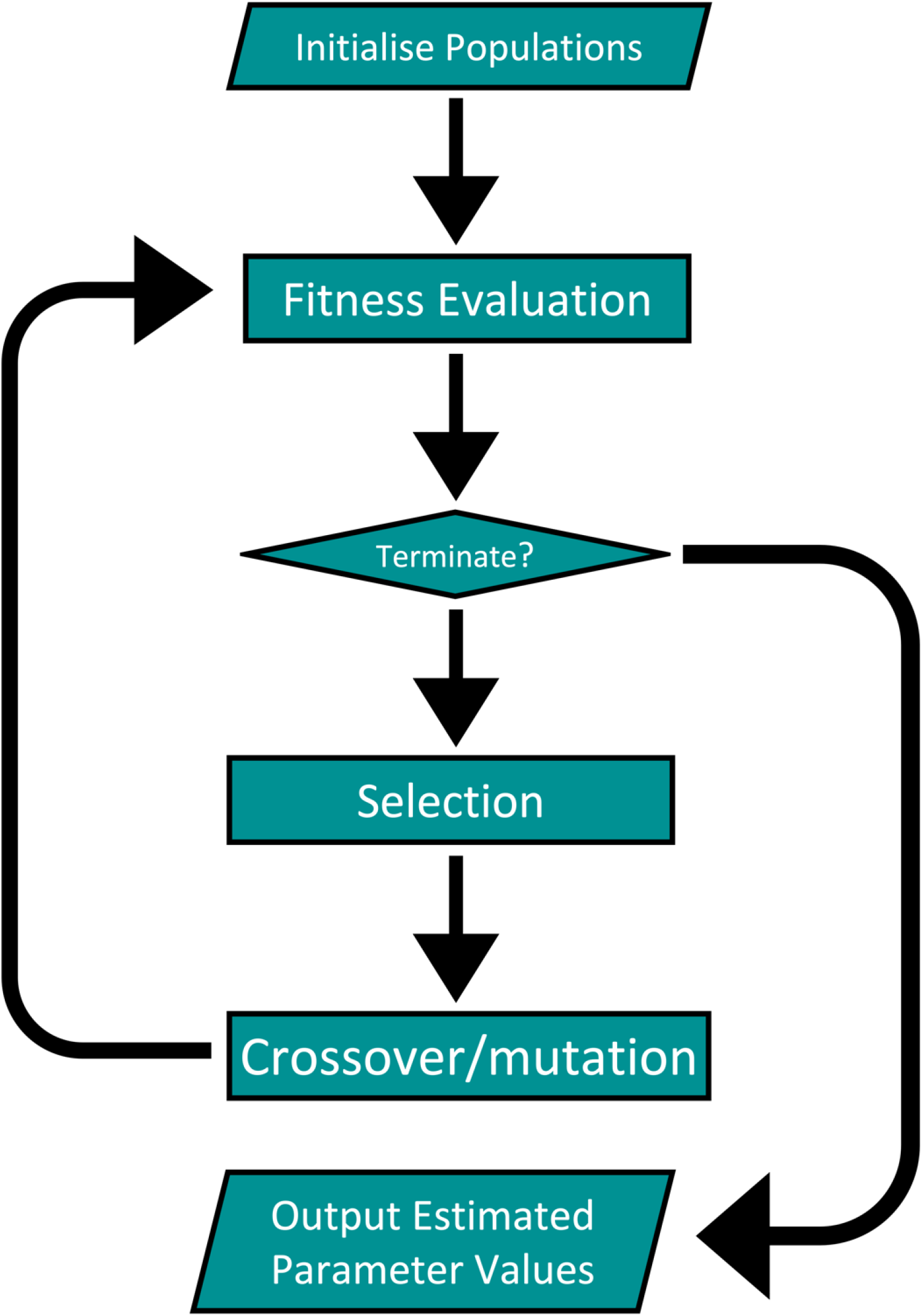
Diagram outlining the processes in the GARS. Each run is populated by an initial 60 parameter sets and continues for 100,000 generations.

## Results

### GARS provides an improved model fit compared to alternative parameter estimation methods

We first assessed the GARS approach to the parameter estimation problem in an established ERK model. This was compared to three other commonly used parameter estimation methods by calculating the Average Relative Error (ARE) between the estimated model solution and the 11 experimental observations across all 11 model nodes^16^.

Parameter values^15^ determined using the EKF method gave an ARE of 20.45% when compared to the experimental data (Table S2), whereas those estimated using Algebraic Difference Equations^16^ achieved an ARE of 12.69%. MATLAB fminsearch optimisation algorithm showed an ARE of 108.27% after 1000 iterations (each parameter value set to the arbitrary value of 0.1) (Table S2). An attempt to modify the parameter values such that they were closer to the true parameter values by setting them to match the final parameter values estimated by EKF increased the ARE significantly to 561.21% (Table S2). As the fminsearch method identifies the local minimum, repeat runs of the algorithm yielded the same solutions. EKF and ADE parameter estimation methods had estimated the parameter values to be between 0 and 1^15,16^. Therefore, the GARS search range was restricted 0 - 1 and showed an ARE of 10.07% (averaged over 5 runs) (Table S2). Despite the previous estimations suggesting that the parameter values should sit in the range 0-1, extending the search range to 0-100 provided a superior model solution with an average ARE 6.97% (over 5 runs) (Table S2). Similar results were obtained using the Heat Shock and Circadian Rhythm models (supplementary tables S1 and S2 respectively).

### GARS provides accurate model fit and prediction in models with noise free synthetic data

Having demonstrated that the GARS provides the most effective method of parameter estimation for model fitting for multiple established biological models we set out to test its robustness to predict unseen data from the Heat Shock Model, ERK Model, Circadian Rhythm Model and NFκB Model.

With 100 noise-free observations, the GARS successfully estimated the parameter values of the heat shock model, which provided an excellent fit for both the given data points and for the unseen true model solution, with an ARE of 0.497% (Figure 3a). Fitting to the circadian rhythm model was very good (Fig. 3b), although with an increased ARE (9.25%) due to the associated oscillatory dynamics. An excellent model fit result was also found when using the larger ERK Model, which gave an ARE of 0.23% (Fig. 3c). For the NFκB model ^17^, 180 noise-free observations over 15 minute TNF stimulation were taken by utilising the parameter values given by Hoffmann et al to define the true model solution. When these observations were used alongside the GARS, an excellent fit to the true model solution was observed, with an ARE of 0.26% (Fig 3d). Similarly, a 30 minutes TNF stimulation also produced a very good prediction of the true solution, providing an ARE of 1.25%.

**Figure 2:**
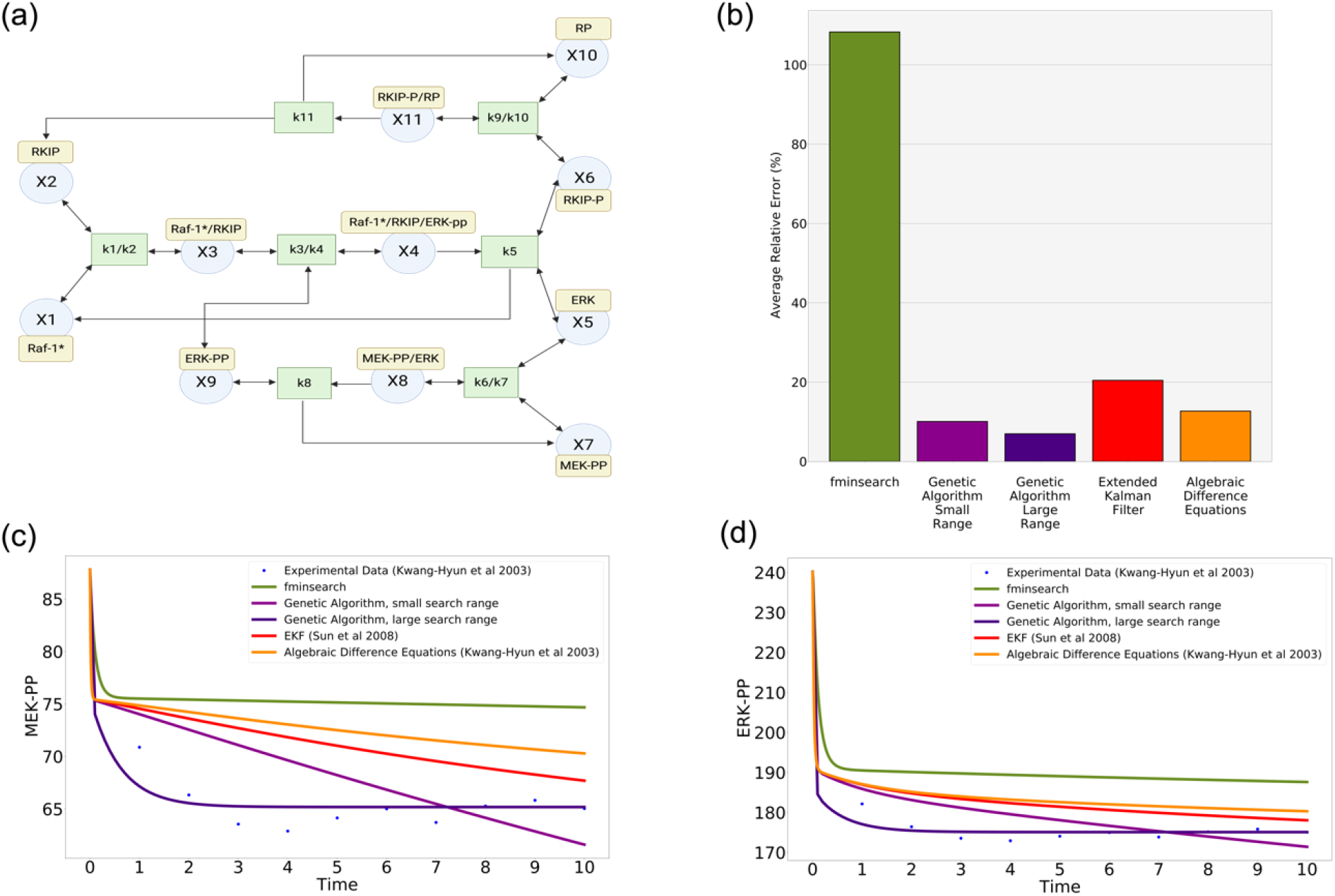
GARS produce a better fit to the data in ERK signalling model: (a) ERK Model developed by Kwang-Hyun et al^16^. (b) Comparison of the Average Relative Errors between the true and estimated model solutions for the 5 methods tested. Standard deviations are unavailable for the Extended Kalman Filter and Algebraic Difference Equations due to the estimated solutions being taken from published literature^15,16^, or for fminsearch due to this method locating only the local minimum. Both GARS were run 5 times, but the standard deviations were too small to be included on this figure (0.001 and 0.002 respectively) and can instead be found in supplementary figure S7. Figure (c)-(d) show the estimated model fitting from each method compared to experimental data^16^ for the ERK-PP and MEK-PP nodes.

**Figure 3:**
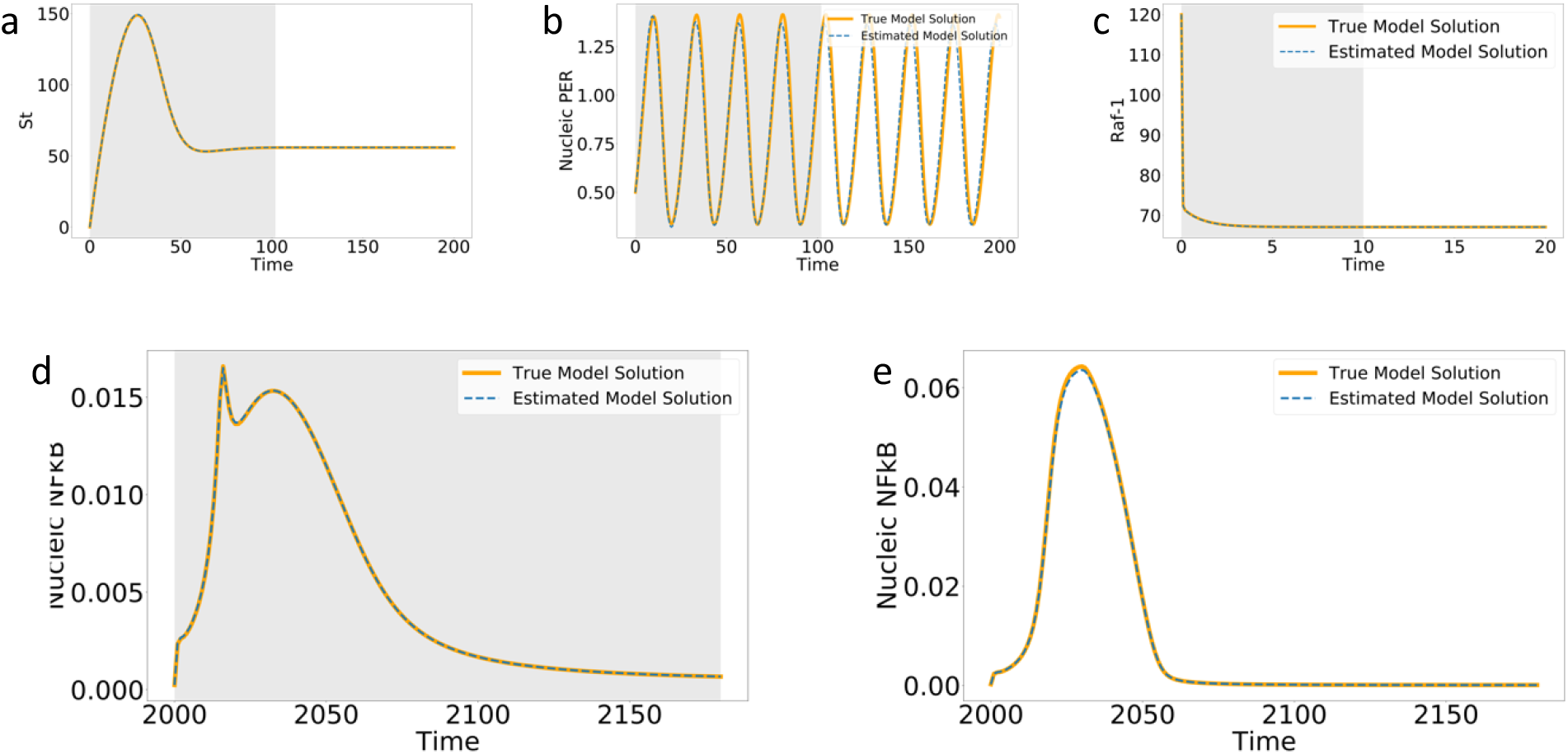
Comparison of true and predicted model solutions when the GARS is provided with noiseless data. Grey background outlines time frame for which the GARS was provided with synthetic observations, whereas a white background indicates that no observations have been taken and the model is predicting system dynamics. (a) Heat shock model^13,18^ with true and predicted solutions. 100 observations were taken over the first 100 time minutes. (b) The Circadian Rhythm model^19^ had observations taken over 100 minutes before the estimated solution from the GARS was able to predict the true model solution for the following 100 minutes. (c) outlines the Raf-1 node of the ERK model with observations taken over the first 10 minutes to provide a true and predicted model solution^16^. (d) shows the NFκB model^17^, with the 15 minute TNF observations being utilised for model fitting, before using the estimated parameter values from the GARS to predict the 30 minute TNF true solution.

### GARS can predict unseen time-course data

To test robustness of the GARS to inherent noise, the Circadian Rhythm model observations (measured over 12 hours per day for 1, 2 or 4 days) plus Gaussian noise with 0.1 variance were used to predict the model dynamics for the next 100 hours after the observations had ceased.

Data from observations of the circadian rhythm over 12 hours (n=23 observations) resulted in a good fit to the estimated model solution (ie less than one oscillatory cycle), but the predictive power of the algorithm dramatically decreased in the following 100 hours and showed significant variation in the replicate solutions (ARE 78.91%, sd 28.27) (Fig. 4a, d). In contrast, over 48hrs (ie two oscillatory cycles, n=24), the GARS produced a good fit of the data over the same time period with good accuracy for both oscillatory frequency and amplitude, but also made accurate predictions over the following 100 hours without observed data (ARE 19.42%, sd 7.88) (Fig. 4b, d). By extending the observations over 96 hours (n=48), the standard deviation of the solutions reduced as compared to 48 hours, as well as providing a more accurate prediction for 100 hours post-observation for both the frequency and amplitude of oscillations (ARE 13.46%, sd 3.72) (Fig. 4c, d).

**Figure 4:**
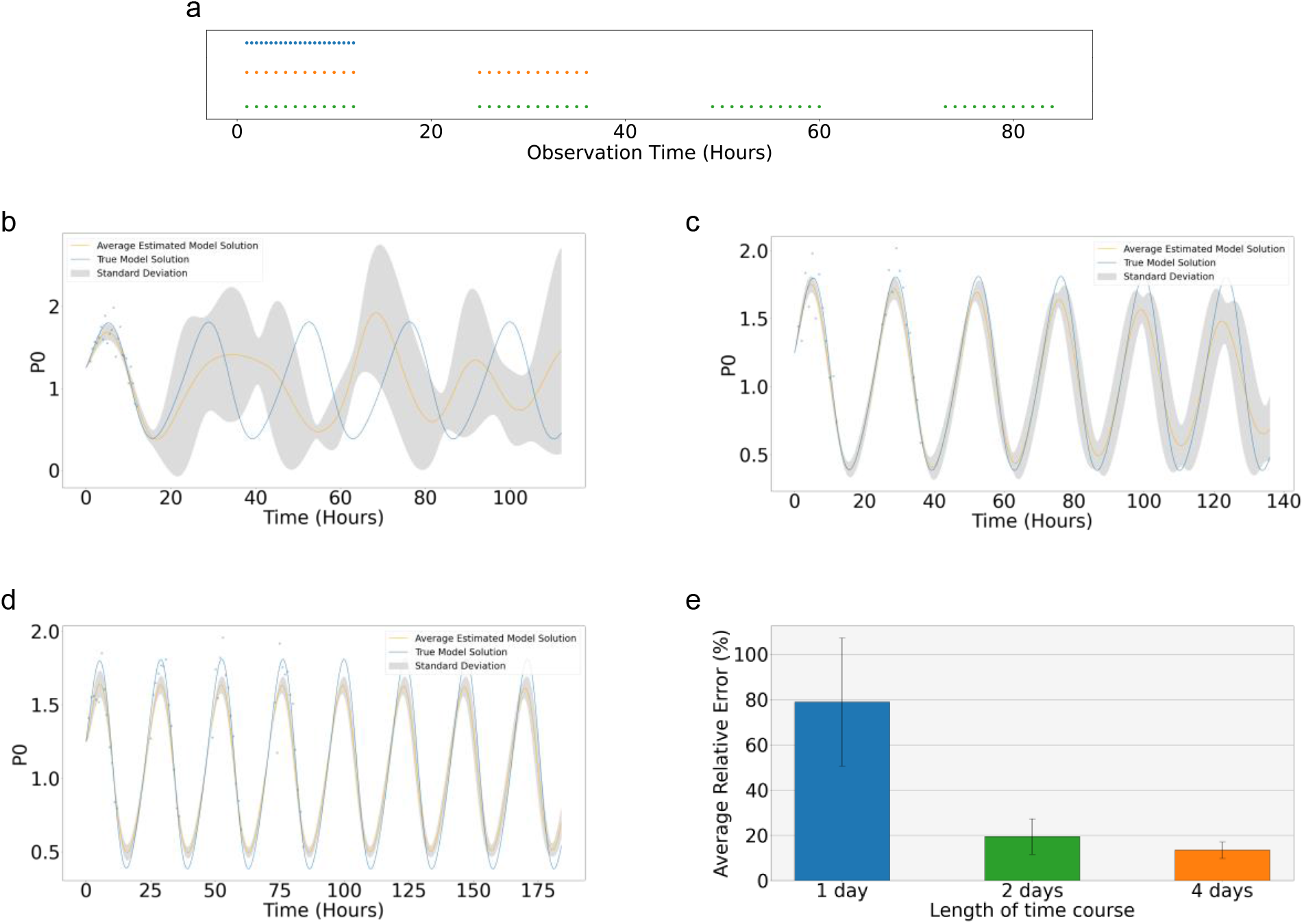
The Circadian Rhythm Model^19^ was used to determine the effectiveness of the GARS when working with realistic data sets. Noisy, irregular observations were taken to represent an average working day in a laboratory setting. (a) Outline of the times at which observations were taken for the 12 hour (blue), 2 day (orange), and 4 day (green) time courses. For the 12 hour time course (b), n= 23 observations were taken with added Gaussian noise of 0.1 variance. The two day time course (c), and 4 day time course (d), consisted of hourly observations over 11 hours in a day, with a 12 hour gap with no observations to represent an overnight break in the data collection, with each having Gaussian noise with variance of 0.1 added to the observations. (e) outlines the reduction in the ARE as the length of the time courses increased.

### Model prediction of unseen biological data from parameters inferred by a GARS

To validate the application of the GARS with biological data, we tested its accuracy to predict over an unobserved period. To achieve this, we examined the NFκB network from fibroblasts stimulated with TNFα as described by Hoffmann et al^17^. TNF stimulation increases nucleic NFκB concentration and these observed data have been previously reported^17^. Using the data from fibroblasts stimulated for 15 minutes, 6 observations were taken over 180 mins and were used to train the GARS.

Figure 5a outlines the model fit to the training data, using the parameter values provided by the GARS. Although the model fit is less accurate than with the synthetic data (ARE 34.6%), it is clear that the algorithm is capturing the underling dynamics of the NFκB system. As we extended the model to predict responses after 30 minutes or 60 minutes of TNF co-culture, we see that it provides model solutions which replicate the system dynamics, with an ARE of 195.8% and 55.16% respectively.

**Figure 5:**
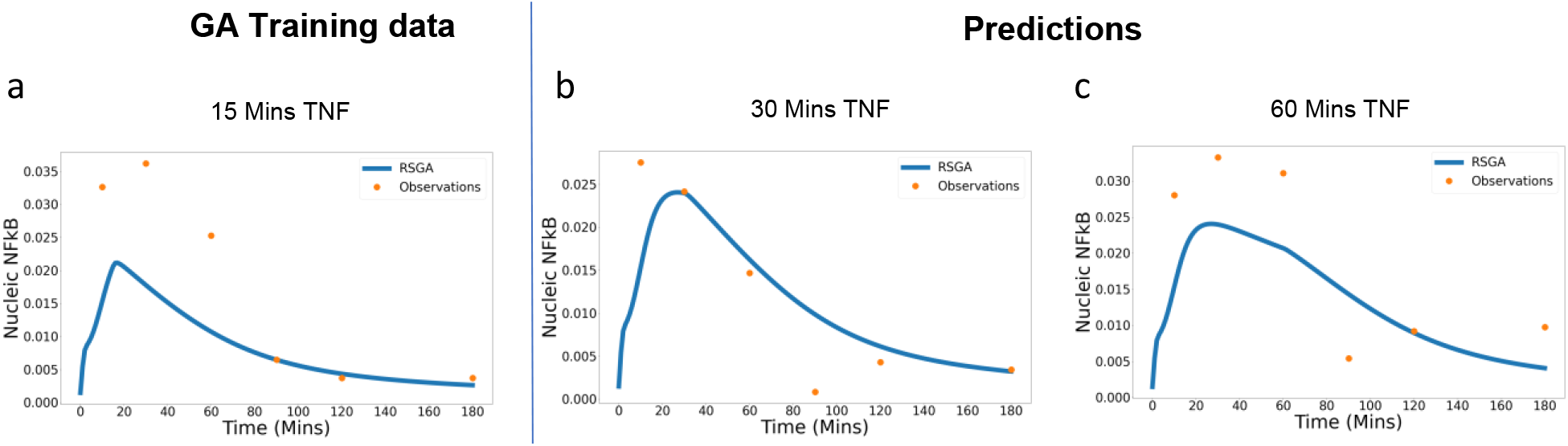
The NFκB model from Hoffmann et al^17^ was used, along with the GARS, to fit to experimental data taken from western blot images after mouse fibroblasts were activated with TNFα for 15 minutes^20^. The estimated parameter values were then used to predict the experimental solutions for the 30 minute and 60 minute TNF activation data sets.

## Discussion

Quantitative models have been developed to describe many biological systems^8,15–17,19,21–23^; however, their predictive capability is limited due to the need for user-defined kinetic rates^1^. These values are often difficult, costly, or even impossible to determine experimentally, and tuning such models by hand can be very time consuming. Several computational techniques have been developed in an attempt to overcome this problem, such as the Extended Kalman Filter (EKF) and MATLAB’s fminsearch function^14,24–26^. However, they also come with their own limitations, such as requiring an accurate initial estimate of parameter values. Additionally, these methods have not thus far been compared on multiple biological networks, nor are they frequently used for the purpose of developing models capable of predicting biological data.

Another limitation of these methods comes from the use of synthetic data for design and validation of algorithms. While convenient and easy to obtain/generate, such syntheses often ignore realistic experimental settings. For example, extended uniform sampling in time is often difficult - either due to difficulty of synchronizing cells during a biological process or simply due to difficulties of a researcher being available in a laboratory at specific points in time -- and measurements are often taken at irregular time points. Additionally, biological and technical noise can influence experimental data, which should be taken into account when generating synthetic observations. Cost of experiments and prior knowledge of how the process responds may also limit the number of time points in which data is acquired. Both these make the possibility of sampling a large number of regularly spaced time points unfeasible.

Rather than an initial estimate of parameter values, as is required by many other parameter estimation approaches, the GARS needs only a search range, and optimal parameter values will be determined within that range. It takes an initial population of parameter value sets and uses them to simulate the chosen model before determining the model fit by calculating the ARE. The GARS then uses ranking selection to determine the parameter value sets which provide the best model fit and performs crossovers and mutations to try and develop an improved solution (Figure 1). This process of selection, fitness evaluation, crossover and mutation is repeated until either a desired fitness is achieved, or the number of iterations reaches a user-defined limit.

We have validated the GARS using realistic synthetic observations to outline the type of data set which could be employed with this method, as well as how we can optimise the algorithm efficiency by planning experiments which take fewer observations over a longer period of time. Additionally, we have established that the GARS can be used as a tool with predictive biological models. By estimating parameter values for an *in silico* model of temporal control executed in NFκB system integrating feedback from IκBβ subunits developed by Hoffmann at al we have been able to accurately predict the effect of perturbing the system with TNF^17^.

Our validation of the GARS demonstrates that it is a tool which can be used with a range of biological models for both synthetic and experimental data, as well as predicting unseen data. Quantitative models generally need to be reductionist in nature due to the computational power needed to simulate them. This means that it is difficult to accurately capture large biological systems using in silico models. Although utilising the GARS can be somewhat computationally heavy, the quantitative results that it produces give it the potential to be a powerful tool for making predictions from biological network models.

Here we have demonstrated that by using a GARS to estimate parameter values for biological network models, we can gain a greater understanding of the underlying model dynamics than with other computational parameter estimation methods. Unlike the EKF and fminsearch methods, the GARS is a parameter estimation tool which requires no pre-existing knowledge of the system kinetic values, and the use of a bounded search range means that parameter values cannot extend beyond their biological range.

By validating the GARS on both synthetic and experimental data, we have shown that this tool can be utilised not only to identify areas of interest or importance within a network, but can also be used to estimate parameters capable of predicting system changes in vitro. By enabling computational models to make accurate predictions about changes to a biological system, we can begin to utilise them as an additional tool to thoroughly investigate such systems, testing variations in ligand concentrations and activation times to determine areas for further experimental investigation, and ultimately infer cellular responses to environmental exposures.

## Methods

### Mathematical Models

#### ERK Network

Kwang-Hyun et al (2003)^16^ first described the system of ordinary differential equations for the ERK network utilised in this study. Figure 2 outlines the model network, and the corresponding equations can be found in supplementary figure S5. The system consists of 11 states which depict the network proteins, along with 11 kinetic parameter values, which were defined using the law of mass action. All decay rates for this model were combined into one term for each equation. Time course data for each of the proteins in the ERK network were generated by Kwang-Hyun et al^16^ and was comprised of 11 observations for each state in the network at equally spaced time points. Where synthetic data was required, 100 observations were taken at regular intervals with no noise added.

Due to the experimental nature of the observed data, no parameter values for this model were assumed to be known. Two search ranges were tested, 0-1 and 0-100. For parameter estimation using the fminsearch algorithm, initial parameter populations were set to the arbitrary values of 0.1 for all parameters. Estimated parameter sets for the EKF and algebraic difference equations (ADE) methods were taken from Sun et al^15^ and Kwang-Hyun et al^16^ respectively.

#### Circadian Rhythm Model

The Circadian Rhythm model used for this study was first developed by Goldbeter in 1995 and consists of 5 equations with 18 parameter values^19^. Synthetic observations from this model for the noise free dataset in figure 3b were taken every hour, with a total of 100 observations. In figure 4 observations were taken every 30 minutes from the 1 hour time point to the 12 hour time point, giving a total of 23 observation points for the 1 day time course, with added gaussian noise with variance of 0.1. The 2 day time course consisted of 24 observations taken hourly from hours 1 – 12, and then from hours 25 – 36. For the 4 day time course, an additional 2 sets of observations from hours 49-60 and hours 73-84 were included. Each observation had added noise using Gaussian random variables with variance 0.1.

#### Heat Shock Model

The Heat shock model from El Samad et al^18^, defines 3 states involved in the heat shock response in E. Coli. The system has 12 parameter values, 6 of which were assumed to be already known during the parameter estimation process and were taken directly from the original model description^18^. The model code was developed in MATLAB following the set of equations in supplementary S3. It was simulated over a timeframe of 200 seconds with intervals of 0.1, from which 200 equally spaced observations were taken to act as known data. Where noisy data was required, Gaussian random variables with variance of 0.1 were added to each of the 200 observation points.

#### NFκB Model

The NFκB model from Hoffmann et al is comprised of 24 states and describes the interactions between IKK, IκB isoforms and NFκB^17^. 5 model parameters were approximated using the parameter estimation algorithms, with the remaining parameters being determined using experimental data and existing literature. Model code was obtained from the BioModels^27^ archive in the form of an SBML file (Model identifier BIOMD0000000140) developed by Harish Dharuri. Experimental data was acquired from the EMSA images captured by Hoffmann et al and analysed using ImageJ to obtain quantitative values for each time point. Where synthetic data was required, observations were taken from the model simulated with parameter values outlined by Hoffmann et al^17^ at 1 minute intervals over a 3 hour time course. Values for experimental observations were generated from western blots^20^ using ImageJ before normalisation.

### Parameter Estimation Techniques

#### MATLAB Fminsearch

Based on the Nelder-Mead Sequential Simplex algorithm^28–30^, the inbuilt fminsearch program from MATLAB^31^ is an optimisation function which can be used to minimise the least squares cost function to find the best fit between the data and the model solution. By minimising the cost function with respect to the parameter values, this method can be used as a parameter estimation technique to find the optimum parameter values to fit the model to the data. The fminsearch method has been used in a number of biological contexts, for example, Kumar et al^32^ used the fminsearch method to estimate the parameters of the JAK/STAT pathway model and Liu et al^13^ implemented it in the Heat Shock model, but concluded that for data with non-Gaussian noise other methods were advantageous.

#### Genetic Algorithm with Rank Selection

Genetic Algorithm with Rank Selections (GARS) are search techniques that allow probing a complex parameter space using a combination of local and large scale perturbations. An analogy is drawn with genetics by viewing local perturbations as mutations and large jumps as cross-over. The key underlying assumption in any genetic algorithm is that the solution is represented (or encoded) in such a way that partial solutions are captured in parts of the representations. This allows retaining good parts of the solution space, and combining partial solutions by cross-over operations. A well-defined fitness function (or inverse of a loss function) needs to be specified and several generations of a population of potential solutions is evolved, retaining good solutions and pruning out poor ones during each generation. Specifically, in the implementations we carried out, the GARS was initialised with a population of 60 parameter sets chosen from a given search range. For each population, the model fit was calculated by determining the Average Relative Error (ARE) between the observations and the estimated model solution, which is subsequently used in the selection stage. Ranking selection was implemented, and the 30 populations with the lowest average relative errors were retained for crossover and mutation. Following this, the chosen populations proceeded to the next generation. The GARS was terminated after 100,000 generations and the optimal solution was taken to be the population with the lowest ARE.

### Computational Simulations

The ERK, Circadian Rhythm and NFκB models and GARS were run in Jupyter Notebook with Python 3.6. Mathematical models were administered using the Tellurium model simulation environment from Choi *et al*^33^. The Heat Shock model was run using MATLAB^31^ and the ODE45 function. All GARS simulations used an initial population size of 60 and were terminated after 100,000 algorithm generations.

### Statistical analysis

The accuracy of the model fit to the data was determined using the ARE. This method allows for unbiased fitting by determining the percentage error between the data and the model at the corresponding time point. The ARE was calculated as described below:

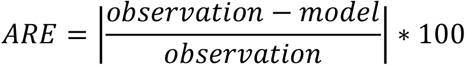

## Acknowledgements

We thank the funding bodies BBSRC and Wellcome Trust, and the iCASE PhD studentship sponsor, Unilever, for providing the resources for this research to take place. We also thank Alexander Hoffmann for his approval for the use of his NFkB Model and corresponding data.

## Author Contributions

Gemma Douilhet - Data curation-Lead, Formal analysis-Lead, Investigation-Lead, Methodology-Lead, Visualization-Lead, Software – Lead, Validation – Lead, Visualisation – Lead, Writing – original draft-Lead, Writing – review & editing-Lead

Mahesan Niranjan – Formal analysis – Supporting, Investigation – Supporting, Methodology – Supporting, Software – Supporting, Supervision – Supporting, Writing-review and editing - Supporting

Andres Vallejo –Formal analysis-Supporting, Methodology-Supporting, Software - Supporting

Kalum Clayton – Formal analysis - Supporting

James Davies – Formal analysis - Supporting

Sofia Sirvent – Formal analysis – Supporting

Jenny Pople - Conceptualization-Lead, Funding acquisition-Supporting, Supervision-Supporting, Formal analysis - Supporting

Michael R Ardern-Jones – Conceptualization-Lead, Formal analysis-Lead, Funding acquisition-Equal, Investigation-Equal, Supervision-Equal, Writing – review & editing-Equal

Marta E Polak - Conceptualization-Lead, Formal analysis-Lead, Funding acquisition-Equal, Investigation-Equal, Supervision-Equal, Writing – review & editing-Equal

## Supplementary Figures

**S1:**
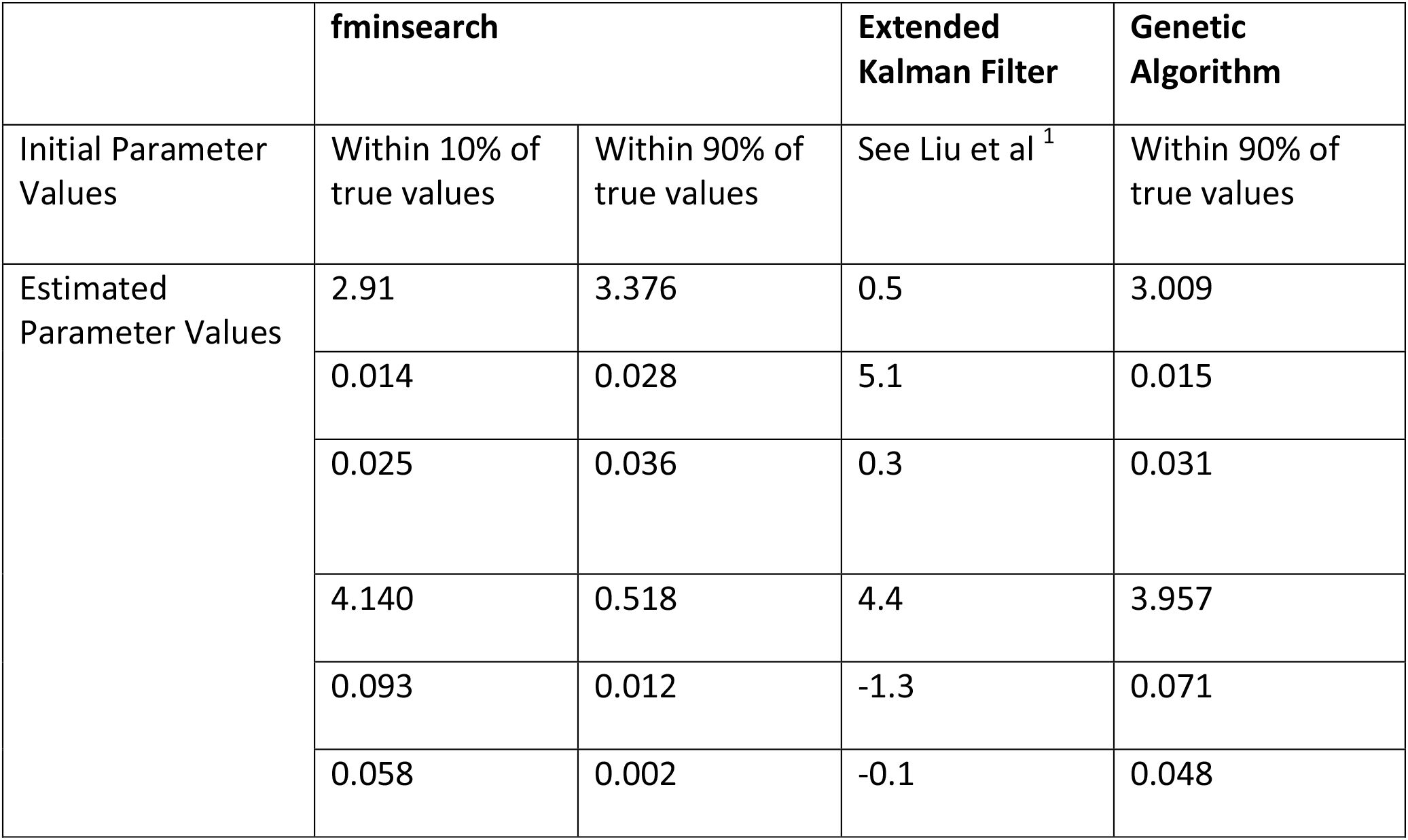
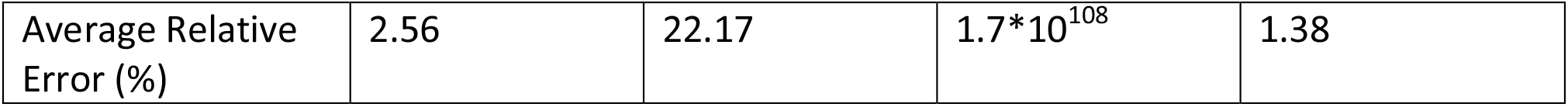
Parameter estimation comparisons for the Heat Shock Model

|  | **fminsearch** |  | **Extended Kalman Filter** | **Genetic Algorithm** |
| --- | --- | --- | --- | --- |
| Initial Parameter Values | Within 10% of true values | Within 90% of true values | See Liu et al <sup>1</sup> | Within 90% of true values |
| Estimated Parameter Values | 2.91 | 3.376 | 0.5 | 3.009 |
|  | 0.014 | 0.028 | 5.1 | 0.015 |
|  | 0.025 | 0.036 | 0.3 | 0.031 |
|  | 4.140 | 0.518 | 4.4 | 3.957 |
|  | 0.093 | 0.012 | -1.3 | 0.071 |
|  | 0.058 | 0.002 | -0.1 | 0.048 |
| Supplementary Figures |  |  |  |  |
| Average Relative Error (%) | 2.56 | 22.17 | $1.7 \times 10^{108}$ | 1.38 |

**S2:**
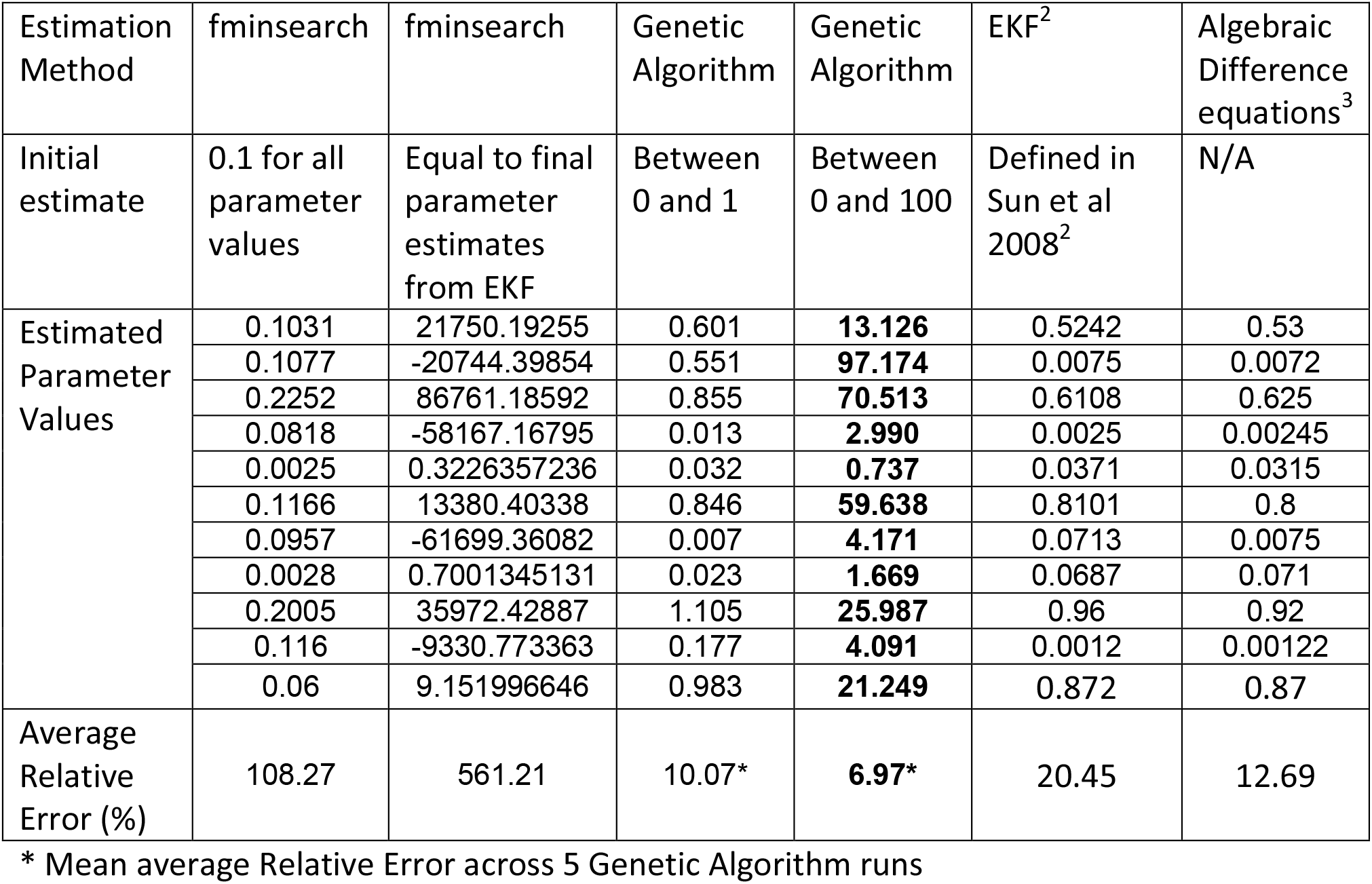
Parameter estimation comparisons for the ERK Model

### S3: Heat shock Model Equations^4^

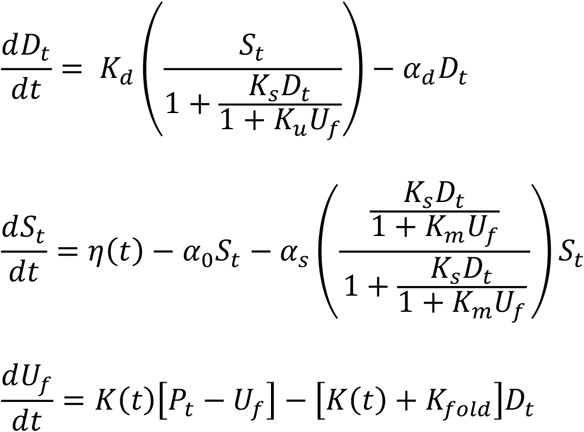

### S4: Circadian Rhythm Model Equations^5^

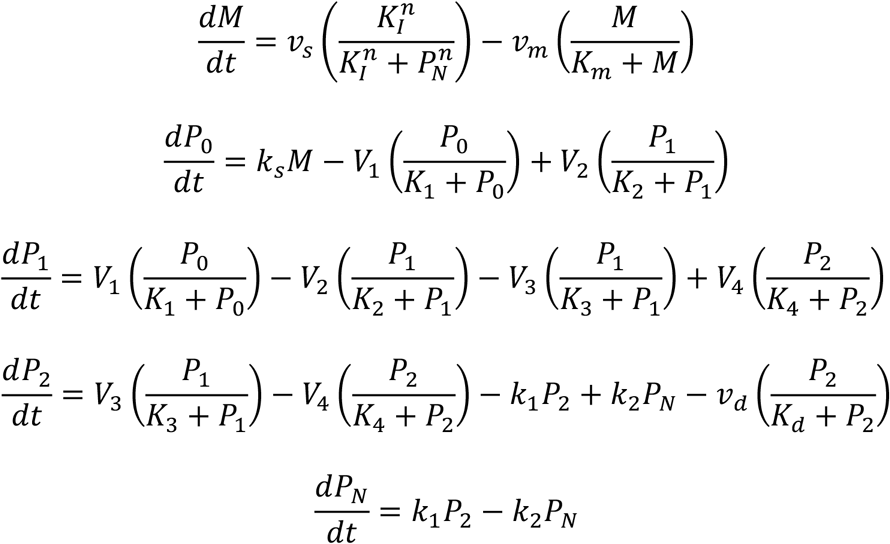

### S5: ERK Network Model^3^

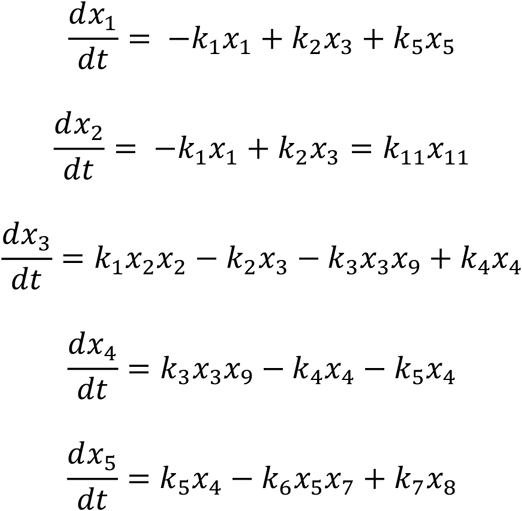

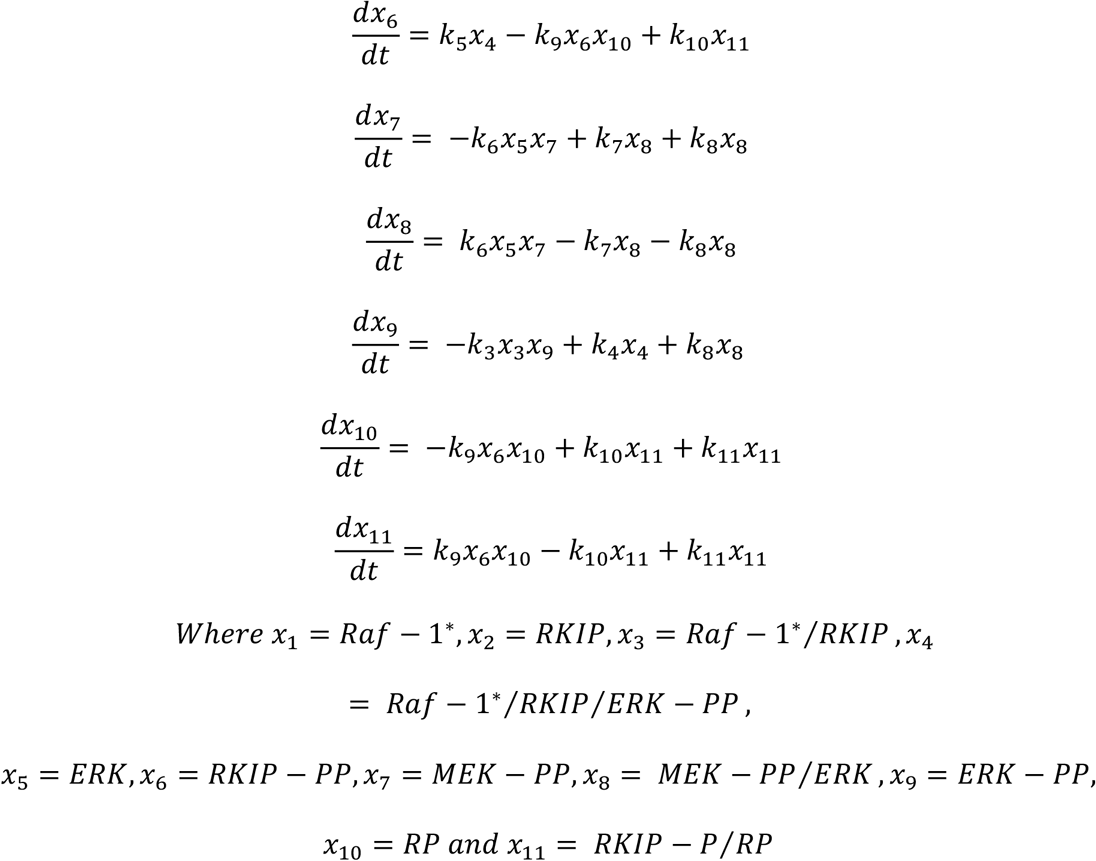

### S6: NFkB Model Equations^6^

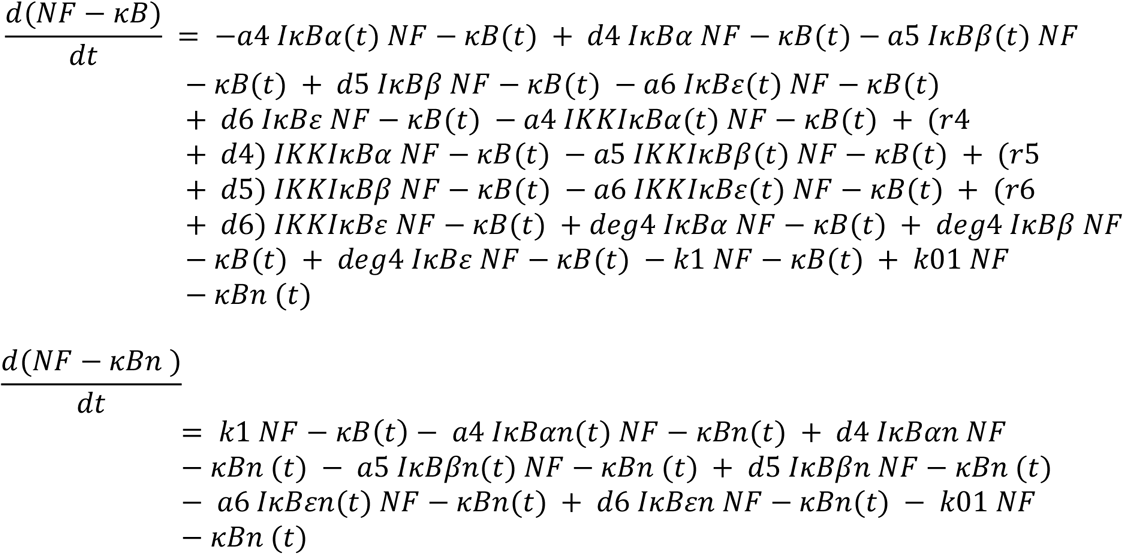

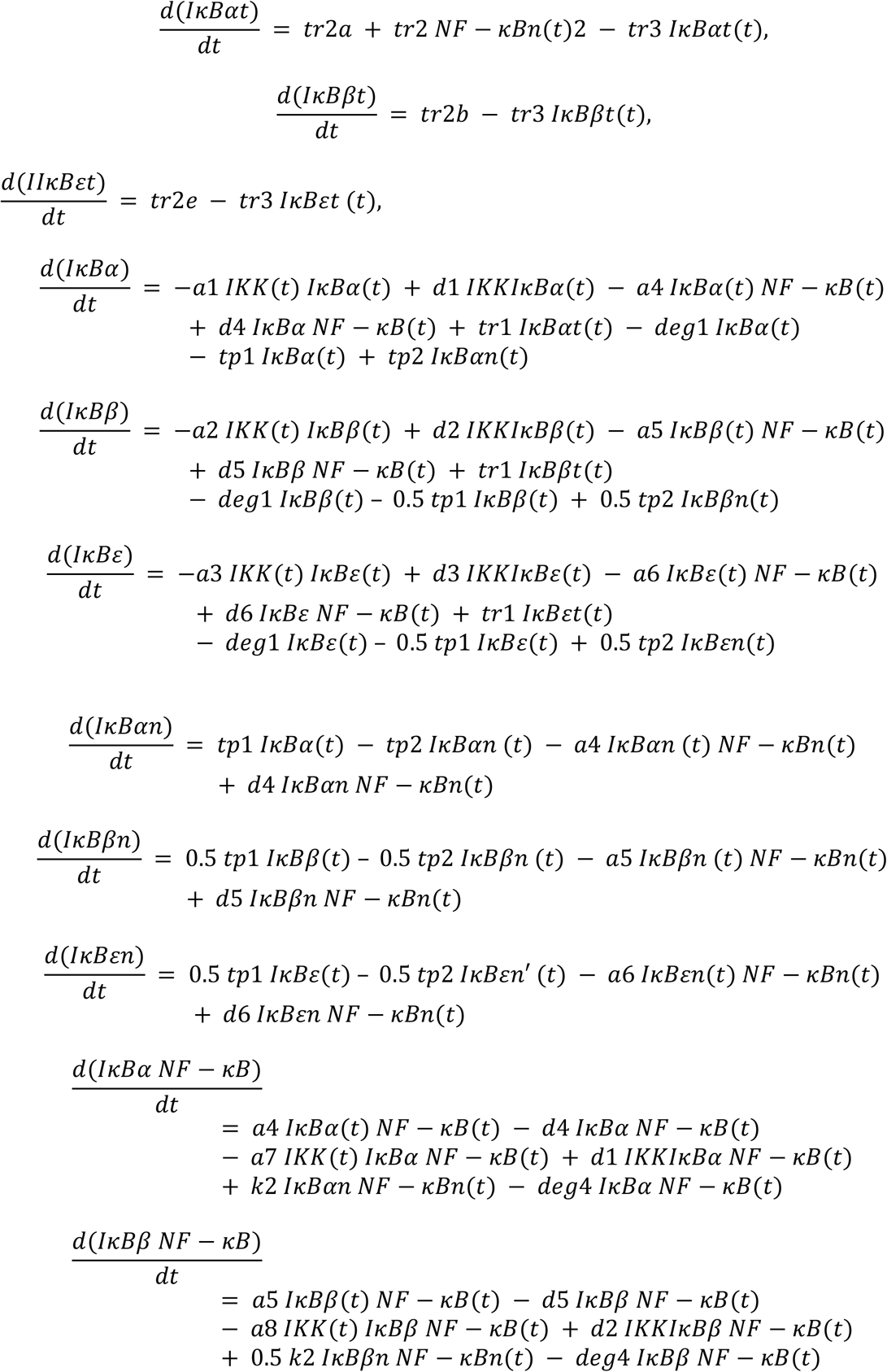

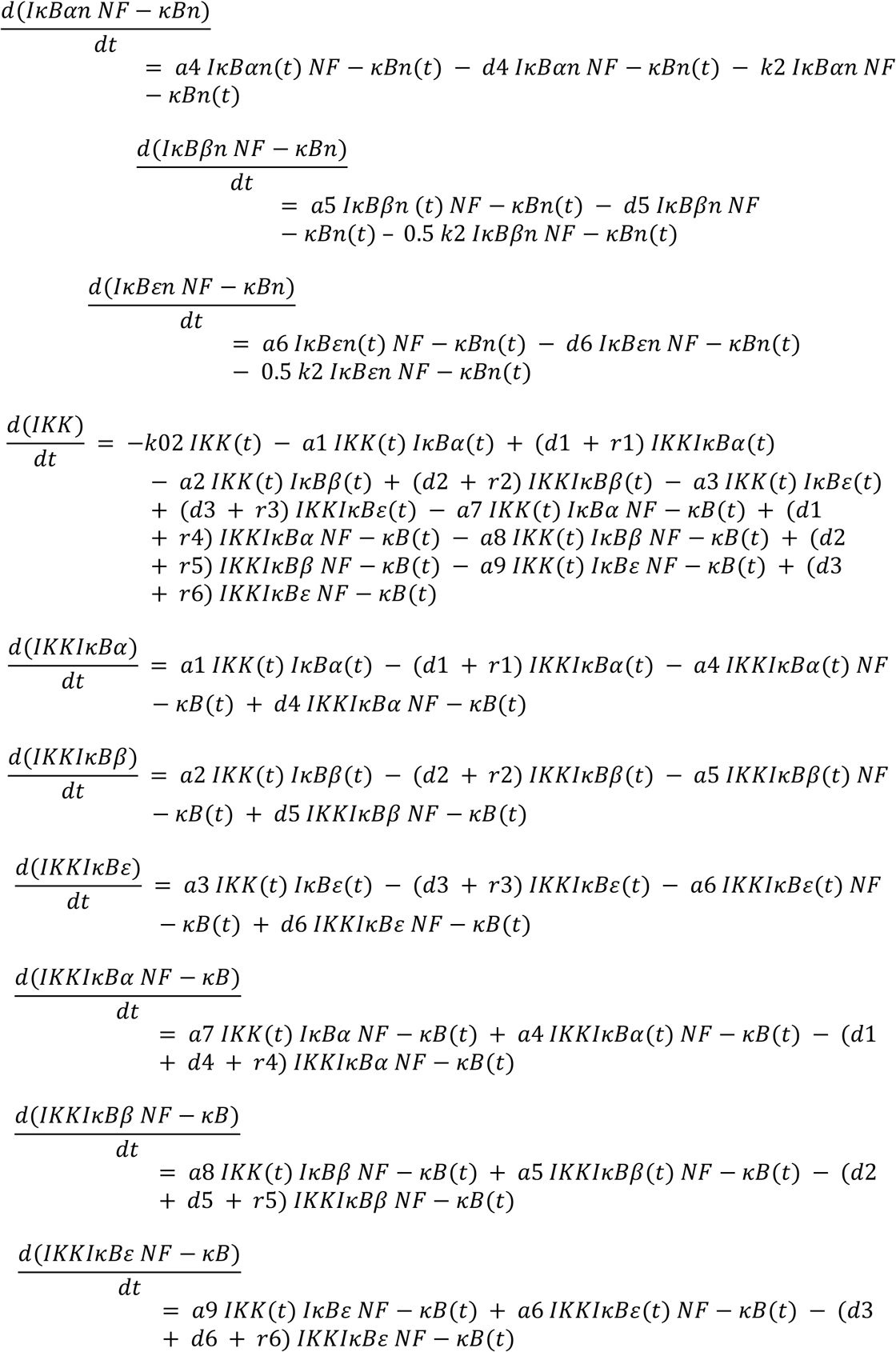

**S7:**
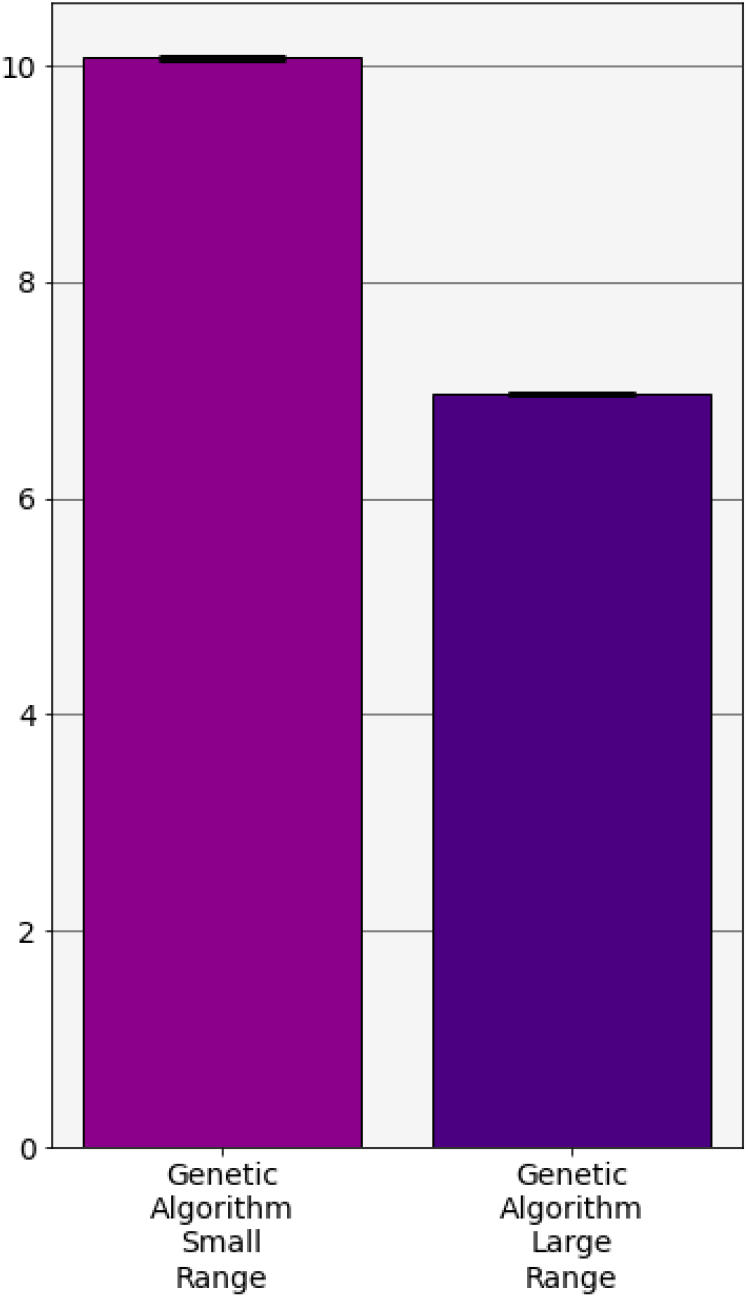
RSGA estimations of ERK model parameter values over multiple runs. Average relative errors using RSGA estimations on the ERK model. Each RSGA was repeated 5 times to give standard deviations of 0.001 and 0.002 for the small and large search ranges respectively.

